# Classifying directed and diffusive transport in short, noisy single-molecule trajectories with wMSD

**DOI:** 10.1101/2022.10.25.513659

**Authors:** Noémie Danné, Zhiqing Zhang, Erwin J. G. Peterman

## Abstract

Fluorescence imaging in combination with single-particle tracking analysis has emerged as a powerful tool to study and characterize the motion of proteins moving in biological media. One of the main challenges in this approach is to reliably distinguish between directed and diffusive transport, especially for short and often noisy trajectories showing distinct, time- and place-dependent modes of motility. In this contribution, we present a windowed Mean-Square Displacements classifier (wMSDc) that is able to reliably (i) identify periods of diffusive and directed transport, (ii) extract position-dependent diffusion coefficients and velocities, and (iii) identify the location of switches in direction or motility mode in short (< 50 time points) and noisy single-molecule trajectories. We compare the performance of this approach to a Hidden Markov Model (HMM) method and a Moment Scaling Spectrum based method (DC-MSS) previously published and show that, in most cases, its performance is superior. We present a wide range of applications: from the movement of whole organisms and cells to protein-DNA interactions *in vitro* and motor-protein dynamics *in vivo*.

**Statement of Significance:** Extracting quantitative parameters from single-particle trajectories is a challenging task, especially in biological samples, which often show a high degree of heterogeneity. Trajectories can reveal switches between different types of motion; directed, diffusive and sub-diffusive motion. Usually, the length of these trajectories and their localization precision are limited by the experimental conditions. Here, we present a novel approach to analyse single molecule trajectories, windowed Mean-Square Displacement classifier (wMSDc) to reliably distinguish between directed and diffusive transport in the short trajectories with a finite precision of localization and integration time typically obtained when imaging single fluorescent proteins in living cells or organisms. We show that, using simulated and a wide range of experimental trajectories, wMSDc is a reliable method to extract motility parameters such as diffusion coefficient and velocity.

## Introduction

Proteins and other biomolecules can be visualized and localized in biological media with sub-diffraction localization precision (*PL* ≪ *λ*/2) (1–5) using fluorescent labeling and imaging (6) in combination with single-particle tracking (7). Using state-of-the-art fluorescence cameras (EMCCD (8), sCMOS (9)) access can be obtained to the dynamics underlying biological processes at video rate. Over the last decade, important focus has been put on developing methods to extract quantitative information from single-particle trajectories (10–14). However, the analysis of short (< 50 time points) and noisy trajectories remains challenging. In single-molecule imaging, such trajectories are commonly obtained because of limiting imaging conditions: pixilation, excess or readout noise of the detector, out-of-focus fluorescence background, undesired autofluorescence from the sample, photo-bleaching/blinking of fluorescent proteins (15) and movement of the biomolecules of interest out of focus. These limitations affect both the localization precision (16) and the length of the trajectory. The challenge in analyzing such trajectories becomes even larger when molecules switch quickly between periods of directed and diffusive transport. In our previous work on intracellular transport in the chemosensory phasmid cilia of *Caenorhabditis elegans* (17–19) we have frequently observed switches between diffusion and directed motion. Switches can be immediate or can involve pauses of variable duration between transport in opposite directions (20). The complex motion of ciliary components in *C. elegans* cilia reveals a wide heterogeneity not only in terms of motion diversity (directed motion and diffusion / sub-diffusion) but also in terms of location-dependent velocity along the length of the cilium (21–23).

Several methods have been developed before to distinguish directed and diffusive transport in single-particle trajectories (12–14, 24–34). Mean-Square Displacement (MSD) analysis has been the most extensively used to analyze the motion of particles and proteins in crowded environments (14, 24–26, 32, 34). This method is relatively straightforward to apply and uses the anomalous exponent of the MSD to differentiate between different modes of transport. However, when the MSD is calculated using a temporal window this limits the temporal resolution and can affect the classification of the motion. Another classification approach makes use of probabilistic models like Hidden Markov Models (HMM). Classification is obtained by estimating the most probable mode of transport for each step in the trajectory based on the states of the previous steps (12, 35–37). However, such probabilistic approaches can result in misclassifications when the of the different modes of transport within a time step are similar and the noise is substantial. Moreover, the number of states that can be considered is limited. In a third approach, Divide-and-Conquer Moment Scaling Spectrum analysis (DC-MSS) the different modes of transport are classified after performing a track segmentation. This method uses a short time window (11 frames) during the initial segmentation allowing to minimize the potential effect of noise. It allows classification of a wide diversity of motion: immobile, confined, free diffusive and directed motion. However, the classification performance is limited by the number of time points in each segment because the MSS exponent values overlap (13). Recently, efforts have been made to develop deep-learning algorithms and combining them with already existing methodologies. The anomalous exponent of the MSD is used either as an input to feed a back-propagation neuronal network (38), or as an output of a Random Forest machine learning algorithm (39). These methodologies are promising but still have some limitations. For example, when the anomalous exponent is used as an input, the temporal window used can affect the classification.

In the current contribution, we propose a windowed Mean-Square Displacement classifier (wMSDc) approach to analyze this kind of trajectories – short, noisy and with velocities and diffusion coefficients that can be location dependent. The MSD is calculated for each point in the trajectory using a time-window previously determined with the trajectory’s characteristics: time between two consecutive points (Δ*t*), localization precision (*PL*) and estimated averaged velocity (〈*v*〉). The classification between periods of directed and diffusive transport is based on the anomalous exponent value (*α*) of the time lag (*τ*) (*MSD* = 2*Γ*. *τ*^*α*^, with *Γ*, the generalized transport coefficient). *α* values are determined analytically for each time points. In order to perform the classification, we use an *α*-threshold value (*α*_*ht*_) determined automatically for each individual trajectory. This paper addresses the following questions: (i) how can *α* be determined analytically?; (ii) what is the optimal time window?; (iii) what is the optimal *α*-threshold value to classify directed and diffusive transport?; and (iv) how can location-dependent velocities and diffusion coefficient be determined? We also compare the efficiency of the classification between wMSDc and two other methods: a HMM based method (12) and a Moment Scaling Spectrum based (DC-MSS) method (13), and discuss the limitations of each approach. Finally, we show several applications of our approach.

## Results

### Windowed Mean-Square Displacement classifier (wMSDc) and ***α*** values: effect of localization precision, integration time, velocity and diffusion coefficient

Based on the anomalous diffusion model, the MSD of a one-dimensional trajectory can be expressed as the product of a constant, called the generalized transport coefficient *Γ*, and a power, *α*, of the time lag

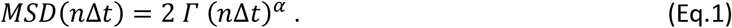

In (Eq.1), Δ*t* is the time between two frames and *n* is the number of frames. In case that a molecule undergoes pure directed transport, *α* = 2, and the average velocity *v* can be determined using *v* = (2 *Γ*)^1/2^ (units in [*m*. *s*^−1^]). In the case that a molecule diffuses freely, *α* = 1. The diffusion coefficient *D* of the molecule then equals *Γ* (units in [*m*^2^. *s*^−1^]). The case 1 < *α* < 2 corresponds to a mixture of directed and diffusive transport occurring within the time-window. For 0 < *α* < 1, MSD scales sub-linearly with the time lag and the displacements of the molecule are limited by local crowding or compartment boundaries (24, 33). Sub-diffusion can also occur when the molecule (intermittently) links to a fixed structure. The case *α* < 0, can occur when a molecule comes back to the initial position using the same path within the time window. As previously reported in the literature, *α* values can be used to distinguish between directed and diffusive transport within trajectories (14, 32). Commonly, *α* is extracted by fitting the MSD curve using a two-parameter fit (*Γ* and *α*). To limit the uncertainties due to the number of free parameters, we choose to calculate *α* analytically using the following equation (40):

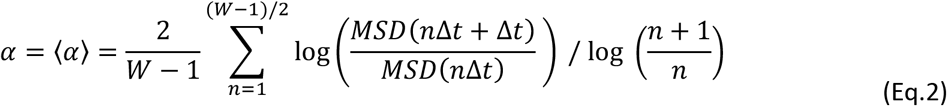

We assume here that the parameter *Γ* is constant and equal to an average transport coefficient determined on the time scale of the sliding window. To increase the accuracy, we average *α* over the first (*W* − 1)/2 time points of the window where *W* is the width of the sliding window. Each *α*_*i*_ value calculated within each window *W*_*i*_ (a sliding window centred around the *i*^th^ data point in the trajectory) is addressed for all the time points included in *W*_*i*_ as explained in Figure S1. In this way, each point has one or more *α*_*i*_ values. The final *α* value for each point corresponds to the average of all the *α*_*i*_ values (〈*α*〉). Figure 1A and 1B show examples of this wMSDc analysis of simulated trajectories for either directed or diffusive transport, respectively. The averaged *α* values obtained were equal to 1.75 and 0.81 for directed and diffusive transport respectively. These values are smaller than the theoretical values (2 and 1) because the localization precision adds random noise to the position of the molecules (see below).

**Figure 1:**
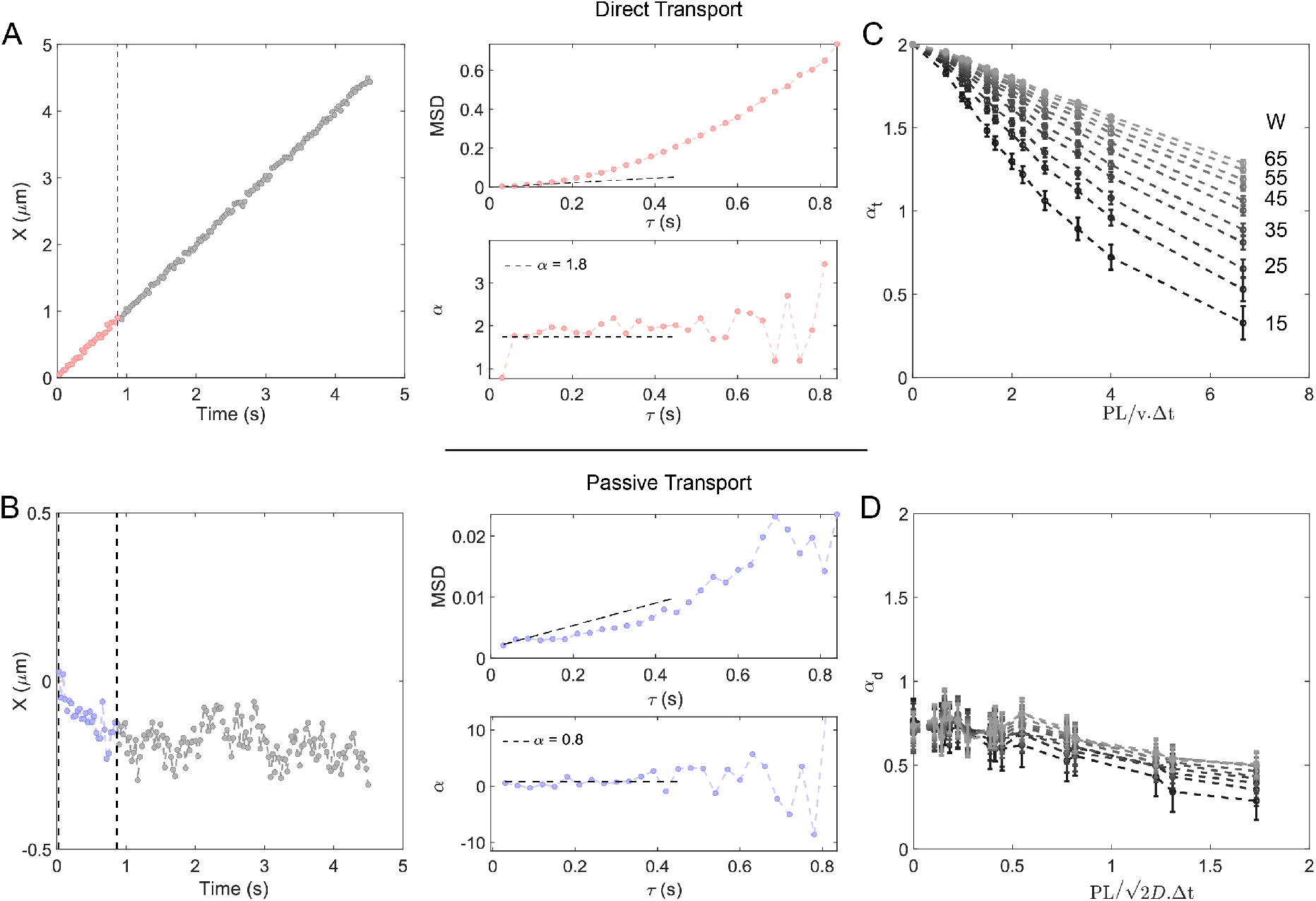
MSD analysis and α values during directed and diffusive transport. A) Simulated trajectory of a single particle undergoing directed transport (*v* = 1 μm. s^−1^, Δ*t* = 30 ms and *PL* = 30 nm); MSD (top right) and α value (bottom right) extracted using a sliding window width *W* of 31 frames. The dashed dark lines show the linear fit of the 3 first lag time for MSD and the averaged α value over the *W*/2 first lag time. B) Simulated trajectory of a single particle undergoing diffusion (*D* = 0.01 μm^2^. s^−1^, Δ*t* = 30 ms and *PL* = 30 nm); MSD (top right) and *α* value (bottom right) extracted using a sliding window width *W* of 31 frames. The dashed dark lines show the linear fit of the 3 first lag time for MSD and the averaged *α* value over the W/2 first lag time. C) *α* values determined for directed transport as a function of the ratio *PL*/(*v* ∙ Δ*t*) using different sliding Window widths (*W*). D) *α* values determined for passive transport as a function of the ratio *PL*/(2*D* ∙ Δ*t*)^1/2^ using different sliding Window widths (*W*). Velocities (*v*), diffusion coefficients (*D*), integration times (Δ*t*) and Localization precision (*PL*) values used for the graphs C and D are reported in Supplementary Table 1.

In order to understand to what extend this discrepancy with the theoretical *α* values is due to localization precision (*PL*), integration time (Δ*t*), velocity (*v*), diffusion coefficient (*D*), and window width (*W*) used, we generated trajectories using different velocities (*v*: 0.3 – 1 μm. s^−1^), diffusion coefficients (*D*: 0.01 – 1.8 μm^2^. s^−1^), integration times (Δ*t*: 30, 60, 100 ms) and localization precision (*PL*: 0, 30, 60, 90 nm), reflecting typical values we have obtained in single-molecule trajectories of ciliary components in the chemosensory cilia of *C. elegans* (20–22). For each data set, we simulated 20 XY-trajectories of 200 steps (see parameters in Supplemental Table 1). In the X-direction, *in-silico* molecules undergo unidirectional directed transport and, in the Y-direction, the molecules diffuse freely. The X and Y-directions are analyzed separately using Eq.1 and 2. As reported before (12), alpha values change as a function of the window width. For this reason, the window width used to calculate alpha varies between 15 frames and 65 frames. The minimal window width is set to 15 frames to limit the fluctuations of *α* between consecutives time points (see Figure S2). Figure 1C shows that *α* values of directed transport (*α*_*t*_) decrease when the ratio *PL*/(*v* ∙ Δ*t*) increases. The decrease of *α*_*t*_ is less pronounced when *W* (expressed in number of frames in Figure 1) increases. Figure 1D shows that *α* values of diffusing molecules (*α*_*d*_) decrease with increasing ratio *PL*/(2*D*. Δ*t*)^1/2^. (Note that *v* ∙ Δ*t* and (2*D*. Δ*t*)^1/2^ are equal to the average displacement of a molecule between two consecutive time points for directed and diffusive transport, respectively). For diffusion, the window width does not show a significant effect on *α*_*d*_ values. This could be explained by the fact that the average displacement < X >_*W*_ = 0 during free diffusion (Brownian motion), independently of *W*. In the range of *PL*/(2*D*. Δ*t*)^1/2^ values tested, *α*_*d*_ is always below 1. Taken together, *α*_*t*_ values for directed transport depend strongly on the ratio *PL*/(*v* ∙ Δ*t*) and the window width *W* used to calculate them, while *α*_*d*_ values calculated for diffusive trajectories are always below 1, showing a small dependency on the ratio *PL*/(*v* ∙ Δ*t*) and the window width *W*. These simulations have shown clear directions to make a proper choice of *W* based on the values of *α*_*t*_ and the characteristics of the individual trajectories.

### Classification between diffusive and directed transport

To classify directed and diffusive transport in single trajectories using wMSDc, we need to set an appropriate window width (*W*) as well as an appropriate *α*-threshold value (*α*_*Th*_). Since the calculation of *α* values depends on *W*, the *optimal* couple (*W*, *α*_*Th*_) needs to be set to perform the classification.

#### Window width and *α* threshold values (*W*, *α*_*Th*_) for classification

In this section, we describe the three steps used to determine the appropriate minimal window width and threshold (*W*, *α*_*Th*_). The first step is to determine the appropriate window width (*W*). *W* should be as small as possible, in order to detect and classify ‘stretches’ with the highest possible time resolution. Here, we define a stretch as a continuous part of a trajectory in which the mode of motility (directed motion or diffusion) is constant. For efficient and reliable classification, the difference (*α*_*t*_(*v*, *PL*, Δ*t*) − *α*_*d*_(*D*, *PL*, Δ*t*)) between *α*_*t*_ for directed motion and *α*_*d*_ for diffusion needs to be maximized. This is because, in the final step, an *α*-threshold, *α*_*th*_, with a value between *α*_*t*_(*v*, *PL*, Δ*t*) and *α*_*d*_(*D*, *PL*, Δ*t*) will be used for classification. A larger difference between *α*_*t*_(*v*, *PL*, Δ*t*) and *α*_*d*_(*D*, *PL*, Δ*t*), will result in a classification that is more reliable and less sensitive to noise. In Figure 2 we depict and describe the different substeps to obtain the appropriate minimal window width for different cases. The first substep is to determine the average and the minimal velocities in the set of trajectories (〈*v*〉 and *v*_*min*_) and to calculate the ratios *PL*/(〈*v*〉. Δ*t*) and *PL*/(*v*_*min*_. Δ*t*). The ratios are represented by two vertical red dashed lines in Figure 2. For each trajectory, the time lag Δ*t* is constant and is defined by the experiment. The localization precision *PL* can be estimated from the global MSD (i.e. MSD calculated for the entire trajectory) by extracting the intersection at the origin (time lag, *τ* → 0 *s*). Here we assume that the *PL* is independent of time and location. We have shown in Figures 1C and D that the dependency of *α* on the average displacement within the chosen time window *W* compared to the *PL* is stronger for directed motion than for diffusive motion. Taking this into account, the window width *W* needs to be chosen such that for directed motion at the lowest velocity within a set of trajectories, *v*_*min*_, *α*_*t*_(*v*_*min*_, *PL*, Δ*t*) is larger than 1. This first criterion (Criterion 1,① in Figure 2), is necessary but not sufficient for an efficient and reliable classification because the difference *α*_*t*_(*PL*/(*v*_*min*_ ∙ Δ*t*)) − *α*_*d*_(*D*, *PL*, Δ*t*), for *D* = *D*_012_ is too small (< 0.2) and this can affect the classification accuracy. This effect can be particularly important in the case shown in Figure 2C for which the velocity is constant. In order to maximize the difference *α*_*t*_ − *α*_*d*_ ∀ *D* and *v* (∀ meaning for all values), we set a another criterium on the averaged *α*_*t*_ value obtained for the averaged velocity 〈*v*〉 estimated within the set of trajectories, *α*(*PL*/(〈*v*〉 ∙ Δ*t*)). For this second substep, we use the maximal *α* value that we can obtain when the velocity is minimal, *α*_*W*=65_, to set the lowest *α* value (*α*_*W*_) we want to obtain for the averaged velocity 〈*v*〉. The objective is to maximize the average value of *α*_*t*_, ∀ *v* obtained for the entire trajectory, and minimize its variations, while using the smallest window width possible. This second criterium can be expressed mathematically be the following expressions:

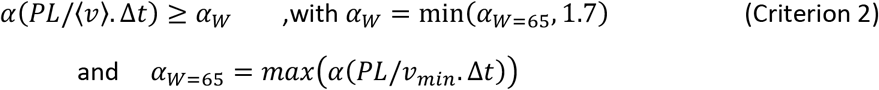

**Figure 2:**
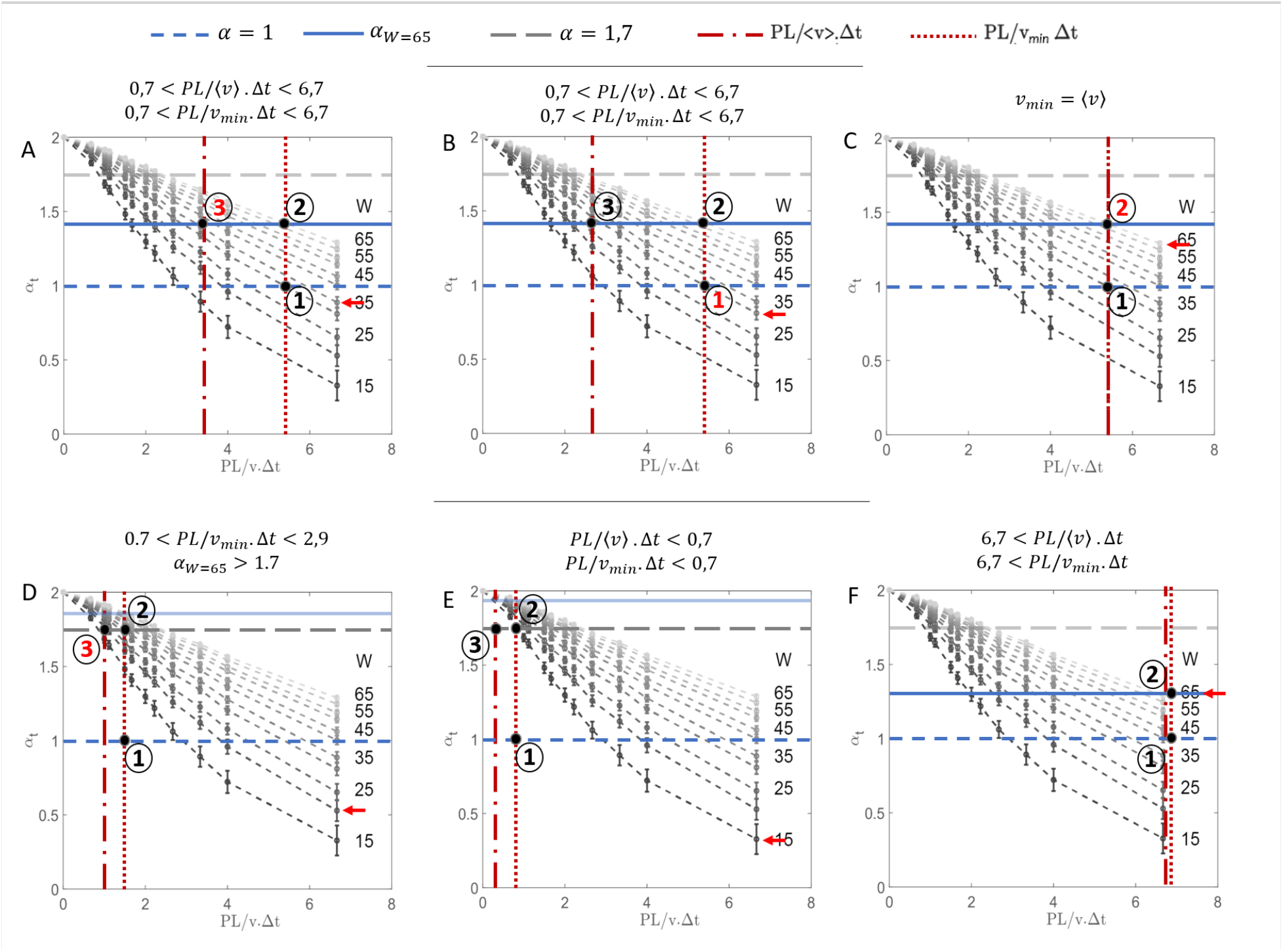
Substeps to choose the appropriate minimal window width *W*. Simulated data are the same as in Figure 1C. The dashed and dotted vertical red lines indicate the coordinates for the ratios *PL*/(〈*v*〉 ∙ Δ*t*) and *PL*/(*v*_*min*_ ∙ Δ*t*). The dashed horizontal blue line corresponds to *α* = 1. The horizontal blue line corresponds to *α* = *α*_*W*=65_ and the dashed grey line to *α* = 1.7. The annotation ① indicates the first criterion (*α*_*t*_(*PL*/(*v*_*min*_ ∙ Δ*t*)) > 1). The annotations ② and ③ indicate the *α*_*W*_ value for the ratios *PL*/(*v*_*min*_ ∙ Δ*t*) and *PL*/(〈*v*〉 ∙ Δ*t*). The red colour for the annotation indicates the most important criterion limiting the value of the minimal appropriate window width. The red arrow indicates the value of the appropriate window width. The different substeps are show for 6 examples. A) *PL*/(〈*v*〉 ∙ Δ*t*) and *PL*/(*v*_*min*_ ∙ Δ*t*) range between 0.7 and 6.7. In this case the second criterion ③ imposes the value of the minimal appropriate window width. B) *PL*/(〈*v*〉 ∙ Δ*t*) and *PL*/(*v*_*min*_ ∙ Δ*t*) range between 0.7 and 6.7. In this case the first criterion ① imposes the value of the minimal appropriate window width. C) *PL*/(〈*v*〉 ∙ Δ*t*) = *PL*/(*v*_*min*_ ∙ Δ*t*). The velocity is constant. This case shows the relevance of the second criteria in order to maximize the difference *α*_*t*_ − *α*_*d*_ ∀ *D* and *v*. D) Particular case for which *PL*/(*v*_*min*_ ∙ Δ*t*) < 2.9 and *α*_*W*=65_ > 1.7. In this case, the lowest value of *α*_*t*_ ∀ *v* (*α*_*W*_) equals 1.7 (dashed grey line). E) Particular case with *PL*/(*v*_*min*_ ∙ Δ*t*) < 0.7. The window width is automatically set to 15 frames. F) Particular case with *PL*/(*v*_*min*_ ∙ Δ*t*) > 6.7. The window width is automatically set to 65 frames.

In Figure 2, *α*_*W*=65_ is represented by a horizontal blue line. In case the ratio *PL*/(*v*_*min*_ ∙ Δ*t*) is above 2.9 (see Figure A-C and F), *α*_*W*=65_ is below 1.7 and thus the lowest *α* value of for the ratio 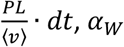, equals *α*_*W*=65_. In the cases of the Figure 2D and E, the ratio *PL*/(*v*_*min*_ ∙ Δ*t*) is below 2.9 and *α*_*W*=65_ is above 1.7. This configuration is not optimal because (i) the curves for the alpha values obtained for the different window widths are near to each other and the determination of the appropriate window width will highly depend on the value taken for the averaged velocity and (ii) a value of 1.7 is high enough to maximize the difference *α*_*t*_ − *α*_*d*_ ∀ *D* and *v*. For this reason, the maximal value of *α*_*W*_ is set to 1.7. The annotations ② and ③ in Figure 2 show the *α*_*W*_ value for the ratios *PL*/(*v*_*min*_ ∙ Δ*t*) and *PL*/(〈*v*〉 ∙ Δ*t*). The third substep is to read from the figure of the simulated data (Figure 2) which window width satisfies the two criteria listed above (red arrows in Figure 2). In the case of Figure 2E and F, the two ratios *PL*/(*v*_*min*_ ∙ Δ*t*) and *PL*/(〈*v*〉 ∙ Δ*t*) are smaller than 0.7 or higher than 6.7. For these cases, the window width is automatically set to 15 or 65 frames respectively. This three-substep strategy has been implemented in our wMSDc approach. In practice, wMSDc determines semi-automatically the window width using the minimal and the averaged velocities (*v*_*min*_ and 〈*v*〉) set by the user. The matrix containing the *α*(*PL*/(〈*v*〉 ∙ Δ*t*)) values for *W* ∈ [15: 1: 65] time points is provided in the Supplemental Material.

After choosing the appropriate window width *W*, the next step is to determine the *α*-threshold value (*α*_*Th*_), used for classification. To test how to do this, we used simulated data based on our experimental data of motor proteins moving bi-directionally in chemosensory phasmid cilia of *C. elegans* (20). These motor-protein trajectories have two types of directional switches: (i) direct turnarounds - switches between directed transport in one direction and in the opposite direction, and (ii) diffusive turnarounds, containing three stretches: directed transport in one direction, diffusive transport (pause), and then directed transport in the opposite direction. In this data, the velocities of motors, duration of pausing and diffusion coefficients while pausing are position dependent. Therefore, we generated data sets using different integration times and localization precision, mimicking the experimental trajectories (see Figure S3 and Methods section for the characteristics of the trajectories). Figure 3A,D show the two types of turnarounds and the *α* values obtained for these two types of trajectories. The colors correspond to *α*_*X*_-values, calculated for the X coordinate using a window width *W* selected by the algorithm (Figure 3B,E - *W* = 15 or 29) as mentioned before. As expected, *α* values during directed transport are above 1, approaching 2 when the localization precision equals 0 (Figure 3E). For a diffusive turnaround (normal case), the threshold, *α*_*Th*_, is determined using the following equation:

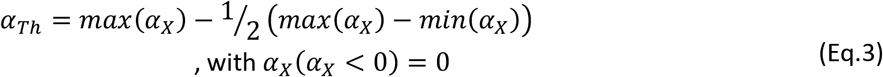

**Figure 3:**
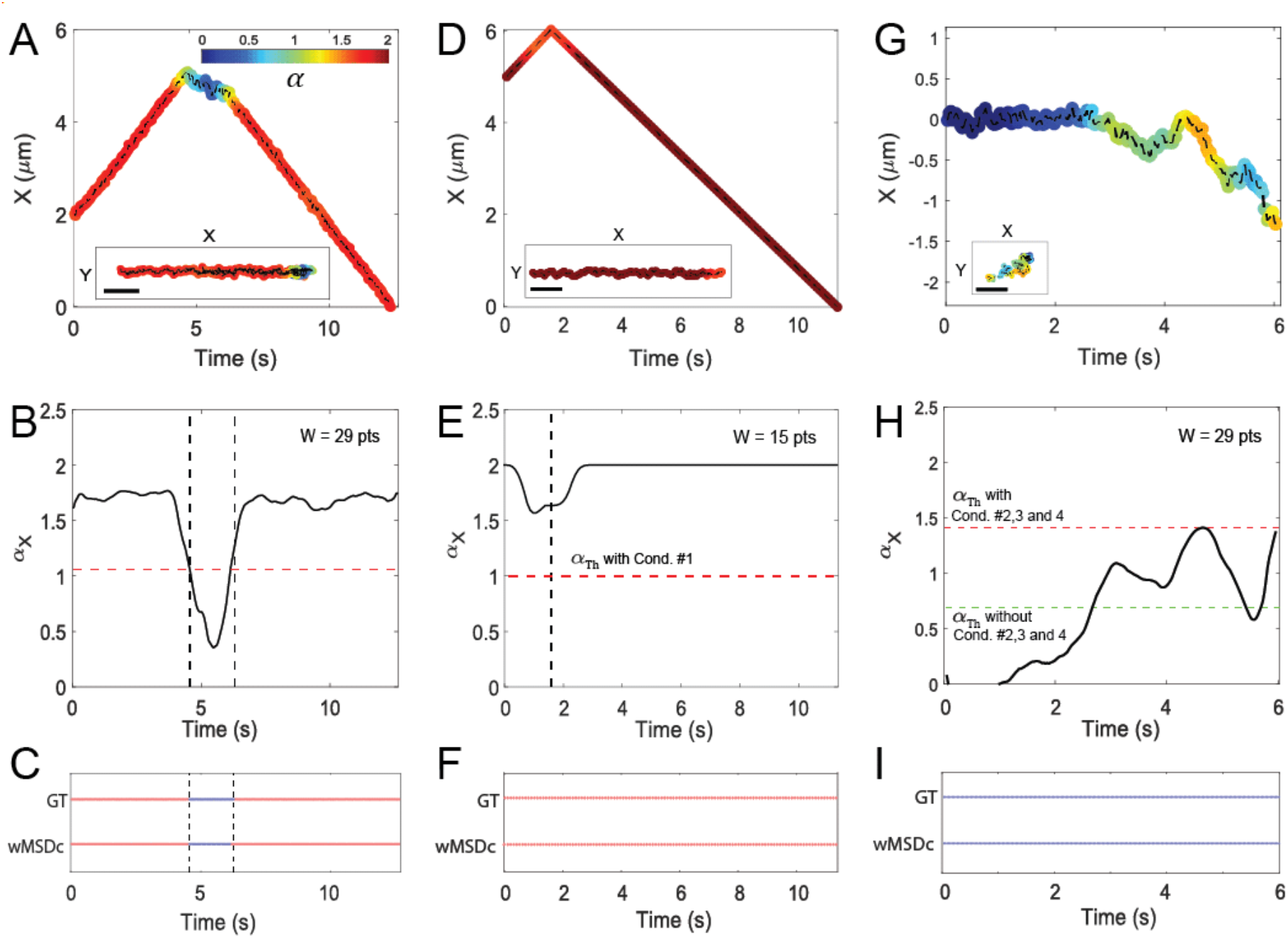
Results obtained with wMSDc after optimization of the Window width and *α*-threshold values. A,D) Examples of simulated trajectories showing a diffusive (Δ*t* = 30 ms and *PL* = 30 nm, *W* = 29 pts) and a direct (Δ*t* = 100 ms and *PL* = 0 nm, *W* = 15 pts) turnaround. The colors indicate *α*_*X*_ values obtained for the X coordinates. B,E) *α*_*X*_ values as a function of time (black line). *α*_*Th*_ values calculated are represented by horizontal red dashed lines. In case of the diffusive turnaround (B), the vertical dashed lines show the transitions between directed and diffusive transport, as injected in the simulation. In case of the direct turnaround (E), the vertical dashed line shows the transitions between the two stretches of directed transport in opposite direction, as injected in the simulation. C,F) Results of the classification within these trajectories using wMSDc. Red: directed transport, blue: diffusive transport. G) Example of a trajectory showing sub-diffusion (0 - 3 s) and free diffusion (3 - 6 s) with Δ*t* = 30 ms, *PL* = 30 nm, D = 0.05 μm^2^.s^−1^ and a radius of 0.1 μm for the confinement. H) *α*_*X*_ values as a function of time (black line). *α*_*Th*_ value calculated using Condition #2 or not are represented by a red dashed line and a green line respectively. I) Results of the classification using wMSDc (Conditions #2. 3 and 4 included). Blue: diffusive transport. GT is Ground Truth (Classification injected in the simulation). Scale bar: 1 μm.

Figure 3C shows that the classification using this equation (wMSDc-row) is in agreement with the Ground Truth (GT-row), and the classification accuracy is 100%.

In case of direct turnarounds (Figure 3E), we note that *α*_*t*_ values are not constant over time despite the fact that we only have directed transport. Indeed, we observe an average decrease of 15% at the turnaround location. In order to classify direct turnarounds as direct turnarounds, we set the following condition (see Figure 3B):

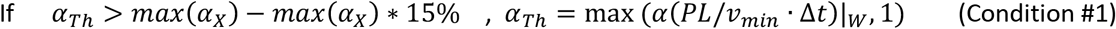

Figure 3E shows that the classification using Eq.3 and Condition #1 (wMSDc-row) within this example trajectory is in agreement with the Ground Truth with an accuracy of 100%. wMSDc detects only directed transport as expected. In order to apply wMSDc to other types of data and not only turnarounds, we also address the specific case where there is no directed transport in the trajectory. An example of a simulated trajectory is shown in Figure 3G. In the first half of the trajectory, the molecule is confined and moves sub-diffusively (*α* < 1, see Methods), and in the second half, the molecule diffuses freely (*α* = 1). Note that even though the molecule diffuses in 2 dimensions (see X-Y trajectory Figure 3G), the *α*_*X*_ value (Figure 3H, at the end of the trajectory) can be larger than 1 (*W* = 29 times points). In this case, the threshold determined with Eq.3 (*α*_*Th*_ = 0.71) is not valid because a part of the stretch showing free diffusion (in this example at times between 2.7 s and 5.5 s, and above 5.8 s) will be classified as directed motion (see Figure 3H, green line). In order to avoid misclassifying such motion as directed transport, we used the *α* values in the perpendicular direction (*α*_*Y*_). Figure S4 shows *α*_*X*_, *α*_*Y*_ as well as *α*_2*D*_ values (the latter corresponding to *α* values calculated using the Mean-Square Displacement in 2 dimensions) for a simulated trajectory containing the following stretches: sub-diffusion, free diffusion, directed motion in one direction, directed motion in the other direction, free diffusion, and directed motion again (in the ‘one’ direction). During diffusive and sub-diffusive motion, the condition max(*α*_*X*_ − *α*_*Y*_) ≤ *α*_*W*_ − 0.5 is almost always true while this is not the case for directed transport. When this Condition #2 is not obeyed, we observed that during diffusive and sub-diffusive motion the Condition #3 *α*_2*D*_((*α*_*X*_ − *α*_*Y*_) = max(*α*_*X*_ − *α*_*Y*_)) ≤ *α*_*X*_((*α*_*X*_ − *α*_*Y*_) = max(*α*_*X*_ − *α*_*Y*_)) − 1/6 and the Condition #4 max(*α*_*Y*_) > *α*_*W*_ − 0.2 are both always true. Based on this observation, we included the Conditions #2, 3 and 4 using *α*_*Y*_ in the algorithm, in order to differentiate, with a higher accuracy, between diffusion and directed motion in trajectories where there is no directed motion. These conditions are based on observations made using the simulated datasets of directed and diffusive transport, and sub-diffusive motion. Figure 3I shows the classification results after the inclusion of Conditions #2, 3 and 4 in the determination of the threshold (dotted red line in Figure 3H). All the time points in the trajectory are classified as diffusive motion, 100% in agreement with the Ground Truth for 90 % of the trajectories (18/20 trajectories). The input parameters are summarized in the Supplemental Table S2. Taken together, wMSDc performs very well in classifying motion in simulated trajectories (in total 290 trajectories divided over 10 datasets, see Supplemental Methods) such as those shown in Figures 3C, F and I, demonstrating the promising power of wMSDc in distinguishing directed motion and diffusion in short and noisy experimental single-molecule trajectories.

#### Comparison of the classification performance of wMSDc, HMM and DC-MSS

In this section we compare the performance of our wMSDc approach in classifying directed and diffusive transport periods in simulated single-molecule trajectories with two other approaches: a Hidden Markov Model (HMM-Bayes) and a method based on Moment Scaling Spectrum (DC-MSS). For the HMM-Bayes method, we used the standard implementation without pooling provided in the reference Monnier et al. (12). In this algorithm, the user selects the number of different states present in the trajectory (K = 2 or 3 states). In order to not bias the results by choosing K = 2 or K = 3, we compared the classification using both values of K (named HMM-2 and HMM-3 in the following). HMM analyzes the trajectories using the X and Y directions and is able to distinguish 2 types of states: Diffusion (denoted D) and a mix between diffusion and directed motion (denoted DV). In order to compare HMM and wMSDc, we consider the DV state in HMM equivalent to the directed motion state in wMSDc. For DC-MSS, we used the algorithm published in the reference Vega et al. (13). DC-MSS also analyzes the trajectories using the X and Y directions and is able to distinguish 4 types of state: immobile, confined, free diffusion and directed motion. In this contribution, we gathered the immobile, confined and free diffusion states together under the class “diffusive transport”.

In order to compare the performance of classification by the different algorithms, we calculated the classification accuracy (%), which corresponds to the number of time points for which the classification is correct (i.e. classifying directed/diffusive transport as directed/diffusive transport), i.e. comparing with the Ground Truth (GT) - divided by the total number of points describing directed/diffusive transport in the trajectory. Figure 4A,B show two examples of classifications obtained for a direct and a diffusive turnaround. For the direct turnaround (Figure 4A, Δ*t* = 100 ms and *PL* = 0 nm), the classification accuracy using DC-MSS, HMM and wMSDc equals 100 %. In case of a diffusive turnaround (Figure 3G, Δ*t* = 30 ms and *PL* = 30 nm), DC-MSS, HMM-2 and HMM-3 classified the entire trajectory as diffusive transport and the classification accuracy was only 14% (57/421), which corresponds to the percentage of diffusive transport in this particular trajectory. In contrast, the classification with wMSDc showed three distinct stretches of different transport modality, with a classification accuracy of 100%. Based on these two examples, we note that the classification of the different types of motion using DC-MSS and HMM is not reliable and depends on the trajectory characteristics (*v*, *D*, *PL*, and Δ*t*), while using wMSDc, the classification accuracy reaches 100%. To investigate the effect of the *PL* and Δ*t* values on the classification accuracy, we repeated this analysis for all the simulated trajectories (*PL* between 0 and 60 nm, and Δ*t* between 30 and 100 ms). We calculated the averaged classification accuracies for directed and diffusive transport in Figure 4C,D. For wMSDc, the classification accuracy for directed transport was relatively constant and above 90% while for DC-MSS and HMM, the results depend strongly on the ratio *PL*/Δ*t*. Their classification accuracy decreases with increasing *PL*/Δ*t*. This effect can be explained by the methods themselves. The classification using HMM is based on the X-Y displacement (Δ*X* − Δ*Y*) distribution and specifically on the center of the distribution in the Δ*X* − Δ*Y* space. When Δ*t* is low, the averaged steps value and thus, the center of the distribution is small. Moreover, when the Noise (*PL*) is high, the width of the distribution is large. The combination of these two effects induces a misclassification of directed transport (GT) as diffusive transport. For DC-MSS, the limitation stems from the number of points in each stretch detected. For stretch durations below 50 time points, the percentage of misclassification increases with increasing ratio *PL*/(*v* ∙ Δ*t*) (13). Directed transport is then misclassified as free diffusion due to an overlap of the moment scaling spectrum values. For diffusive transport (Figure 4D), the classification accuracy depends on the ratio *PL*/Δ*t* for all methods. The classification accuracy for wMSDc is slightly better than for HMM and DC-MSS when the ratio *PL*/Δ*t* is below 0.95, while DC-MSS is slightly better than wMSDc and HMM when the ratio *PL*/Δ*t* is above 1.2 – 1.6. HMM detects diffusive transport with a higher accuracy when the noise is high and when Δ*t* is low as discussed above. The classification accuracy for DC-MSS depends strongly on the number of points detected in each stretch as mentioned above. This effect is important when Δ*t* increases. Indeed, for a fixed pause duration, the number of time points in the stretch is then smaller and the probability of misclassification using DC-MSS increases. For wMSDc, the origin of the misclassification is the width of the window and so in our case is connected to Δ*t*. Indeed, in case of diffusive turnarounds with a pause duration shorter than the window width, *α* values during the pause are higher than 15% of the maximal *α* value. In this case, diffusion is misclassified as directed motion due to the Condition #1 set to determine the *α*-threshold. In conclusion, these results show that wMSDc is more reliable than DC-MSS and HMM to detect directed transport in these short and noisy trajectories. The limitations of DC-MSS and HMM come from the ratio *PL*/(Δ*x*) (with Δ*x* = Δ*X*, Δ*Y*, the displacement between two frames) and the number of points detected in the segment for DC-MSS. For diffusive transport, the limiting factor for wMSDc and DC-MSS is the number of points during the pauses, while for HMM the limiting factor is the ratio *PL*/(Δ*x*).

**Figure 4:**
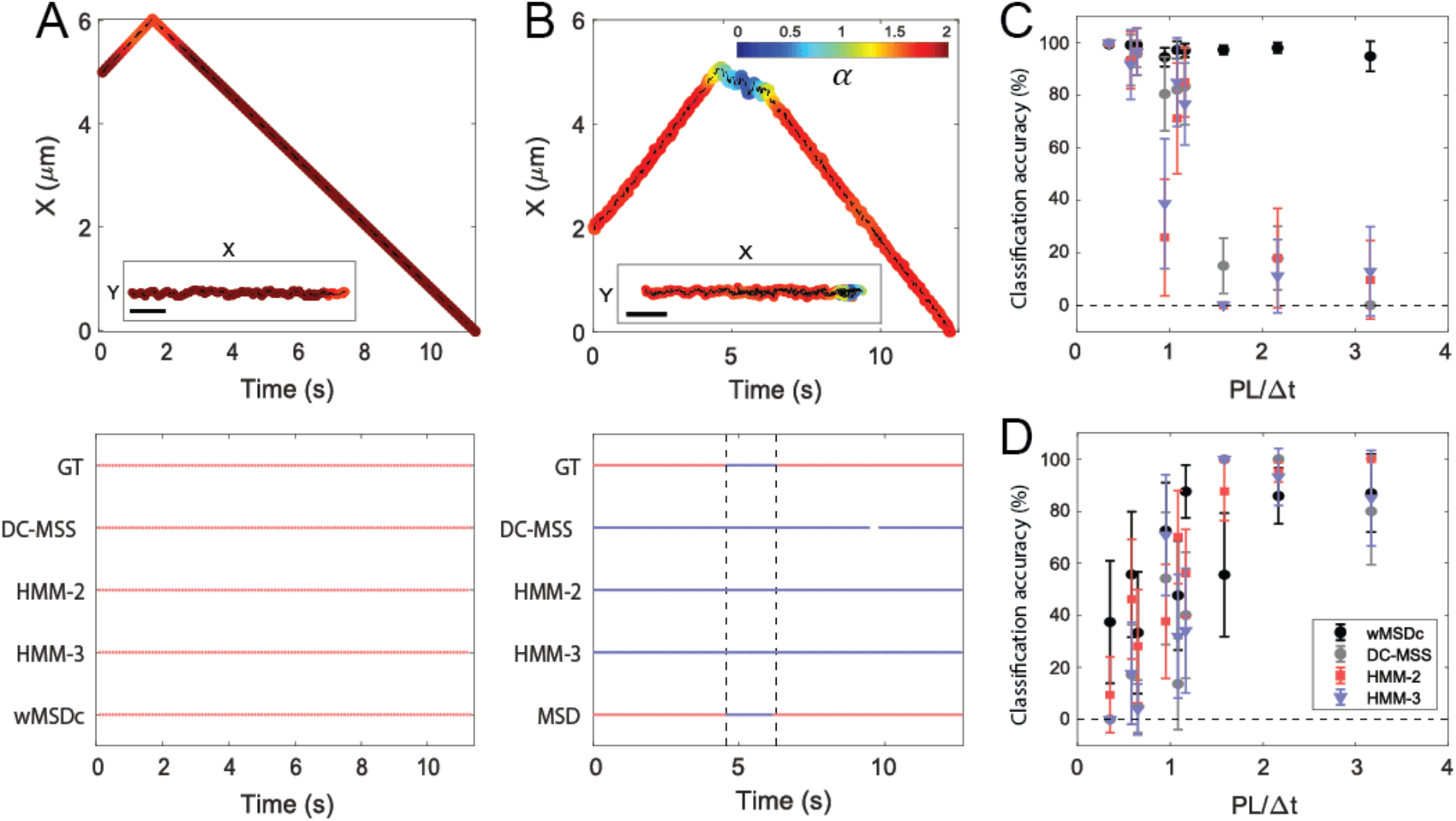
Comparison of the classification accuracy of DC-MSS, HMM and wMSDc as a function of the ratio *PL*/Δ*t*. A,B) Comparison of the classification using the three methods for a direct and a diffusive turnaround. Red: directed transport, Blue: diffusive transport. The dotted vertical lines in B indicate the beginning and end of the diffusive part. GT is Ground Truth, and HMM-2 and HMM-3 indicate the results obtained using HMM with an initial number of states equal to 2 and 3 respectively (see also main text). C) Classification accuracy during directed transport, D) Classification accuracy during diffusive transport. Scale bars: 1 μm.

#### Classification between anterograde and retrograde directed transport

In addition to the classification between directed and diffusive transport, directed transport can be sub-classified based on direction in anterograde and retrograde directed transport. This sub-classification can be important in cases such as the chemosensory cilium of *C. elegans* for which the velocity of transport depends on directionality, since different motor proteins are involved. To distinguish anterograde and retrograde directed transport, we use the sign of the velocity smoothed over several time points (in most case 30, but less (e.g. 15) in case of short stretches, see Applications) to filter the noise coming from the localization precision (sign +: anterograde, sign -: retrograde).

### Velocities and diffusion coefficients: effect of the localization precision and integration time

After we have found a reliable way to classify anterograde directed transport, retrograde directed transport and diffusive transport using wMSDc, the next step is to determine how velocities and diffusion coefficients can be extracted from the data. In order to detect local variations of these parameters, we first used the method published in the work of Holcman et al. (32), where velocities and diffusion coefficients were estimated from the displacement between two consecutive time points *x*(*t*) and *x*(*t* + Δ*t*):

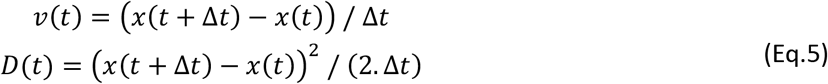

We found, however, that this approach can be substantially biased by the localization precision. To quantify this effect, we used the simulated data presented above. We extracted for each set the averaged values of the velocity or diffusion coefficient and the standard deviation of the mean. The parameters of the simulation can be found in Supplementary Table 1. For directed transport, the averaged velocity calculated with Holcman’s method (Eq.3) was in agreement with the velocity values fed into the simulations (supplementary Table 1, Figure 5A). The standard deviation of the mean of the velocity (*σ*_*v*_) increases linearly when the ratio *PL*/Δ*t* increases (Figure 5C, grey filled circles) as expected. In order to minimize *std*(*v*), our strategy was to use more time points to calculate the velocity. Indeed, when the number of points increases, the standard deviation of the mean decreases (Figure 5C, red circles and Figure S5B). In our method, position-dependent velocities are calculated using a sliding window of 9 time points. Within each time-window, we fit the curve *X*(*n*_*t*_. Δ*t*) = *v*. (*n*_*t*_. Δ*t*) (*n*_*t*_ = 1,. .9 in the sliding window) and extract the velocity. In this way, *σ*_*v*_ is less than 25 % of *v* as shown in Figure 5C (light red line). For diffusive transport (Figure 5B), the averaged diffusion coefficient depends on the number of points used to calculate the diffusion coefficient. The diffusion coefficients calculated using Eq.3 (in grey) differ from the diffusion coefficients fed into the simulations (black dashed line). The standard deviation of the mean of the diffusion coefficient (*σ*_*D*_) depends on the diffusion coefficient (*D*_*th*_), *PL* and Δ*t* as shown in Figure 5D. The ratio *σ*_*D*_/*D*_*th*_ scales, by definition, quadratically with the ratio *PL*/(2*D*_*th*_. Δ*t*)^1/2^. The polynomial coefficients obtained are different depending on the approach used to determine the diffusion coefficient. As expected, it can be clearly seen that the ratio *σ*_*D*_/*D*_*th*_ is smaller and increases less when the diffusion coefficient is calculated using the Mean-Square Displacement and a Window width of 15 frames. Using the Eq.5, the ratio *σ*_*D*_/*D*_*th*_ strongly depends on the ratio *PL*/(2*D*_*th*_. Δ*t*)^1/2^ with standard deviation of the mean, *σ*_*D*_, exceeding 10 times the diffusion coefficient value, *D*_*th*_, in extreme cases. In order to minimize this dependency, we fit the curve (*x*(*t* + *n*_*d*_. Δ*t*) − *x*(*t*))^2^ = 2. *D* * (*n*_*d*_. Δ*t*) + *b*, with *n*_*d*_ = 1, 2 and 3 and extract the diffusion coefficient, *D*. Using this method, the ratio *σ*_*D*_/*D*_*th*_ is reduced with a factor of almost 2 when the level of noise is 1.6 times higher than the average displacement between two consecutive frames (2*D*_*th*_. Δ*t*)^1/2^ (see Figure 5D, blue filled circles). To summarize, we choose to use *n*_*t*_ = 9 and *n*_*d*_ = 3 combined with a linear fit to extract *v* and *D* instead of using the Eq.3 (*n*_*t*_ = *n*_*d*_ = 2), in order to minimize the standard deviation of the means (*σ*_*v*_ and *σ*_*D*_). Using this approach, we calculated the velocities and diffusion coefficients for the two example trajectories in Figure 4 A and B. The results are shown in Figure S5 C and D. Overall, the results obtained with HMM-2, and wMSDc in combination with Eq.3 are in agreement with the Ground Truth values of the position-dependent velocities (average anterograde velocity = 0.68 *μm*/*s* and retrograde = − 0.62 *μm*/*s*). However, we note a slight deviation of the velocity (Figure S5 C,D, in red) compared to the input value (Figure S5 C,D, in dark) at the location of the turnaround. As expected, the sliding window of 9 time points used to extract the velocity results in an underestimation of the absolute velocity at the location of the turnaround. This issue arises from the structure of the algorithm: the velocities and diffusion coefficients are calculated before the complete classification, and in particular the velocities are used to classify between anterograde and retrograde directed transport. To overcome the underestimation of the velocity at the location of the turnaround, one option is to recalculate the velocities and diffusion coefficients after performing the sub-classification between anterograde and retrograde directed transport. In summary, our wMSDc method is more reliable than DC-MSS and HMM-2,3 in distinguishing directed and diffusive transport in simulated data. The classification accuracy depends on the precision of localization and integration time. The velocity and diffusion coefficient values are overall in agreement with the input values independently of the method used, as long as the classification is correct.

**Figure 5:**
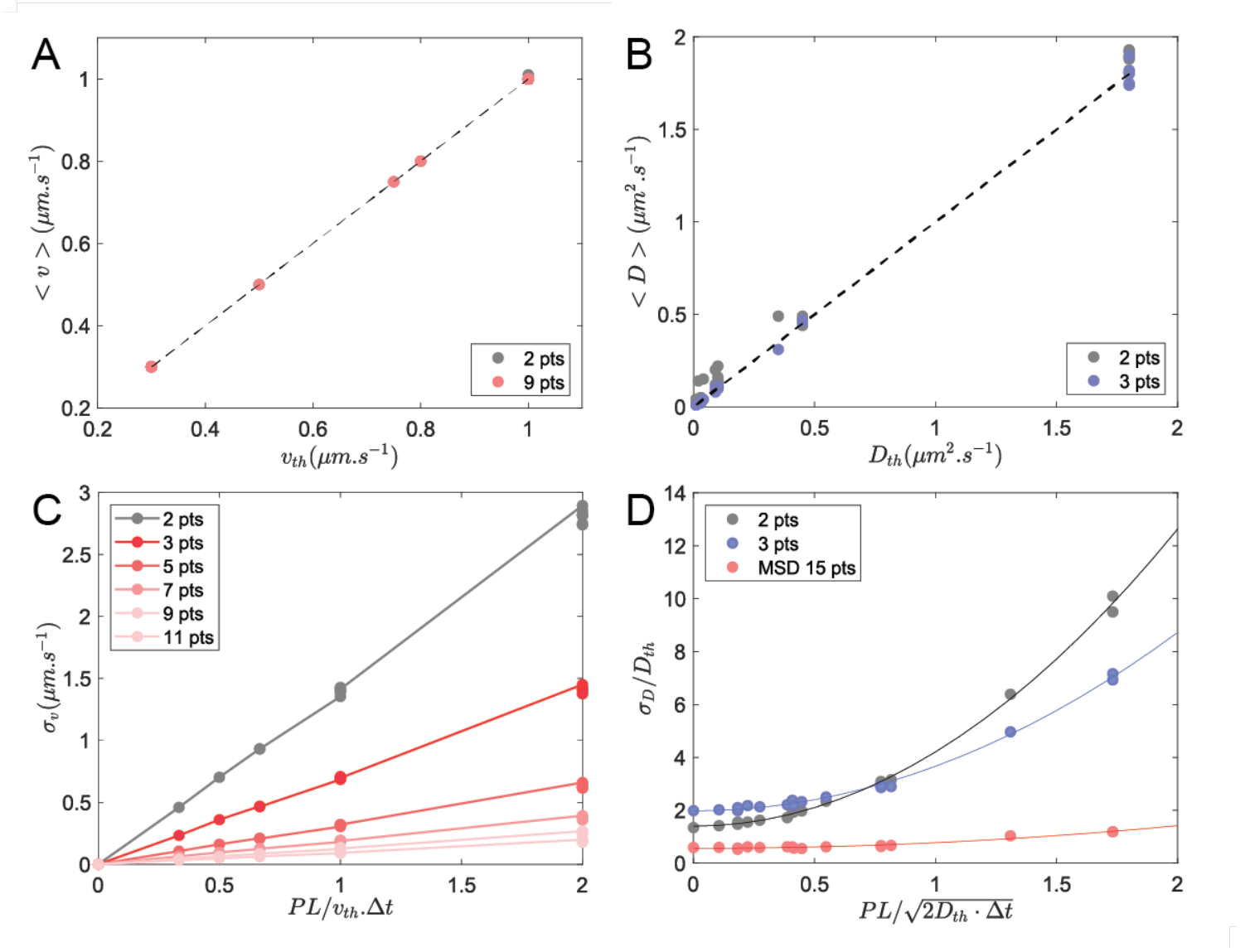
Effect of localization precision (*PL*) and integration time (*Δt*) on the values obtained for the velocity and diffusion coefficient. These results are obtained using the simulated trajectories of Figure 1, with directed motion in the X direction and free diffusion in the Y direction. Values for velocity and diffusion coefficient are listed in Supplementary Table 1 (Set T and Set D). A) Velocity calculated using Eq.3 (between 2 consecutive time points) in grey; Ground Truth velocity shown as black dashed line (see Set T) and velocity calculated using 9 time points in red. Note that the grey and red points overlap because the averaged velocity values obtained using 2 and 9 time points are equal. B) Diffusion coefficient calculated using Eq.3 (between 2 consecutive time points) in grey, Ground Truth D represented as black dashed line (see Set D) and D calculated using 3 consecutive time points in blue. C) Standard deviation of the velocity calculated using different amounts of points. D) Normalized Standard deviation of the diffusion coefficient, *σ*_*D*_/*D*_*th*_, calculated using Eq.5 (Holcman’s method, black solid circles), using our approach (see main text, blue solid circles) and using a linear fit on the Mean-Square Displacement calculated with a window width of 15 time points (red solid circles). The solid lines show quadratic fits.

### Applications

After having validated the performance of wMSDc in classifying motility modes in simulated trajectories, in this section, we apply wMSDc to analyze different kinds of experimental trajectories. Details of the analyses, for example the input average velocities and alpha values are listed in the Supplemental Table S2. We first analyzed trajectories of single motor proteins imaged in the chemosensory cilia of living *C. elegans* (20), similar to the simulated trajectories used to validate wMSDc. In Figure 6A two examples are shown for the classification of the motility in these trajectories in diffusive motion and directed transport. In our analysis, the directed transport is subclassified in anterograde and retrograde transport. Many of these motor trajectories contain directional switches, turnarounds and the subclassification in anterograde and retrograde transport allows classifying turnaround events based on the initial and final motility direction (AR, RA, AA, or RR). Furthermore, the location of the turnaround, whether the turnaround contains a pause or not, what the duration of this pause is, and whether the motor is diffusing during the pause can be extracted. We have applied this wMSDc analysis approach to a complete data set as mentioned above (20).

**Figure 6:**
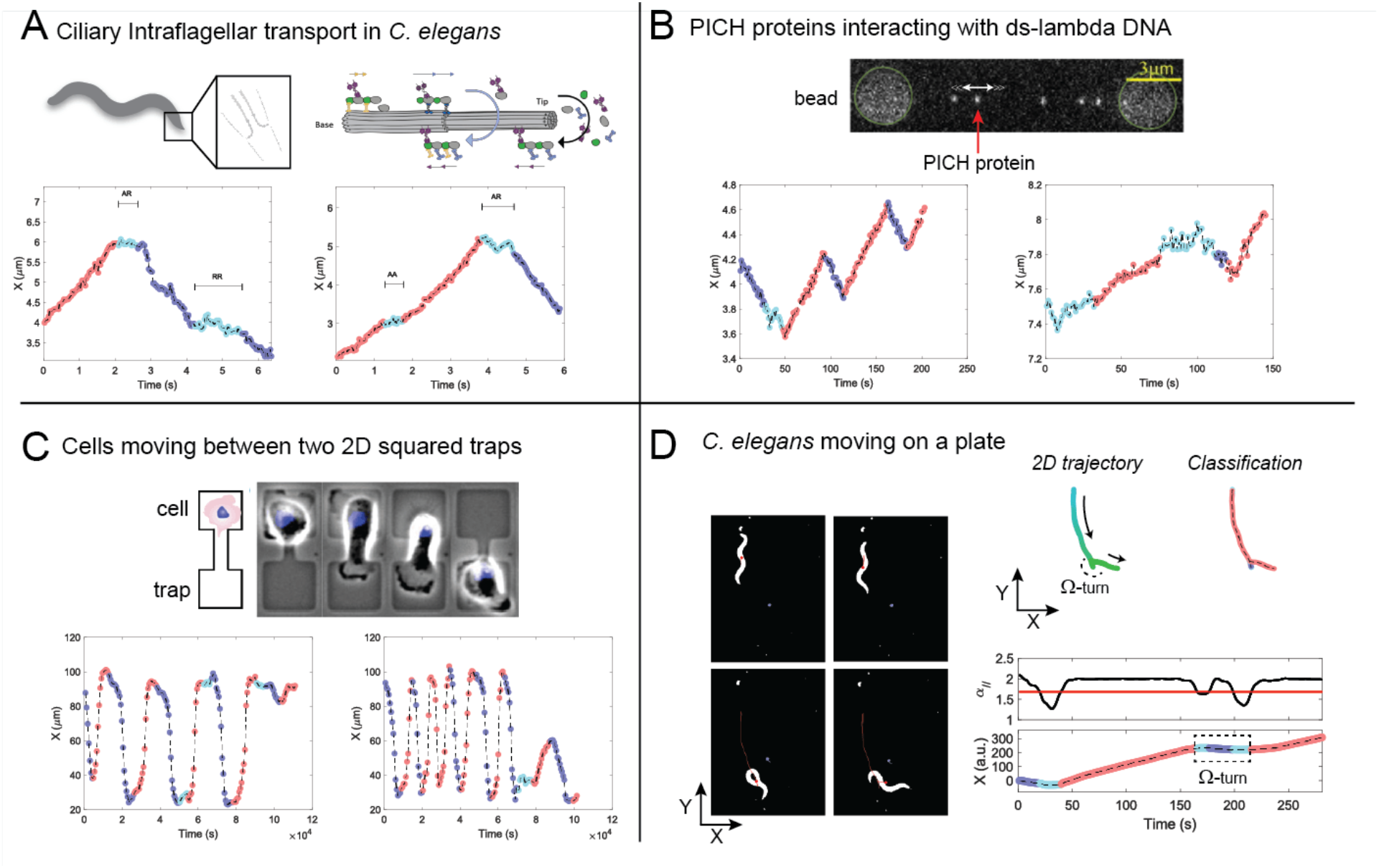
Applications of wMSDc to trajectories from the subcellular to the macroscopic scale: single motor protein trajectories *in vivo,* protein motion *in vitro*, cell motility, and *C. elegans* motion. A) Example of two trajectories of single motor proteins recorded *in vivo* in chemosensory cilia of *C. elegans* (Δ*t* = 30 ms, *PL* = 55 nm and *v* ∈ [0.5 − 1.5] μm. s^−1^) (20). Red: anterograde (A) directed transport, Dark-blue: retrograde (R) directed transport, and Light-Blue: diffusive transport/pauses. AA, AR and RR are classifications of the pauses depending on the type of motion before and after the pause. B) Example of two PICH proteins interacting with a lambda DNA recorded *in vitro* (41). The DNA, with beads attached to both ends is extended and held in place with two optical traps. Δ*t* = 1 – 2.5 s, *PL* = 30 nm and *v* = 10 nm. s^−1^. C) Example of two trajectories showing human cancer cells *in vitro* moving between two 2D-squared traps (42) (37 μm x 37 μm). The position is extracted from the location of the nucleus (in blue). Δ*t* = 10 min, *PL* ≪ averaged steps size and *v* ≈ 1 μm. min^−1^. D) Example of one trajectory showing a *C. elegans* worm performing an Omega-turn (Ω-turn) on an agar plate after touching a repellent-containing liquid droplet (SDS). This typical reaction is a part of the behavioral response of *C. elegans* towards chemical stimuli. Δ*t* = 1 frame, *PL* ≪ averaged steps size and *v* ≈ 2 pxl. frame^−1^.

The second dataset we analyzed using wMSDc involves the protein PICH, a DNA translocase involved in anaphase DNA bridge resolution. We analyzed 1-dimensional trajectories of individual PICH proteins moving along double-stranded lambda phage DNA, extended and held using optical tweezers *in vitro* (41). Similar to the simulated data and the *C. elegans* motor protein data, these trajectories appear to contain turnarounds, the velocities, however, are about two orders of magnitude lower. Since the PICH trajectories are 1-dimensional, we did not apply Condition #2 using *α* values in the perpendicular direction. In Figure 6B, two example trajectories of PICH are shown, using wMSDc. In the left trajectory, several of the fast directional switches (in light blue) are misclassified as diffusion because the window width used in the analysis is larger than the time scale at which these switches occur. This is one of the expected limitations of wMSDc, as discussed above. Apart from this difficulty in correctly classifying frequent directional switches, the overall classification of the PICH trajectories appears sound (Figure 6B), given the low signal-to-noise ratio and the small number of points in the trajectories.

The third dataset we analyzed using wMSDc involves cancerous cells (MDA-MB-231) migrating on a surface between two physical 2D-squared traps (Figure 6C, ref (42)). The authors extracted the trajectories of cell motions by tracking the position of the nucleus as a function of time (shown in blue in Figure 6C, top). To determine the appropriate threshold to analyze these trajectories, we used Eq. 4 to take into account the overall decrease of *α*_*X*_ values induced by fast switches occurring soon after the first switch. Moreover, as the switches occur within a period of time shorter than 30 time points, we adjusted the smoothing used to perform the sub-classification between anterograde and retrograde transport (15 time points instead of 30). The classification results obtained with wMSDc are shown in Figure 6C. Classifications appear sound, demonstrating that wMSDc is able to reliably distinguish directed and diffusive transport for other type of trajectories than the ones we simulated.

The last dataset we analyzed involves 2D-trajectories of C. *elegans* moving on an agar plate before and after touching a droplet of a Sodium deoxycholate (SDS) solution pipetted on the plate, known to act as a repellent. The 2D trajectory of the worm was obtained by determining and tracking the position of the centroid of a segmented image sequence (Figure 6D trajectory in the right upper corner). After touching the SDS droplet, the worm moves backward (in red) and makes a so-called omega-turn (grey-blue, light-blue, grey-blue; between the 160^th^ and 210^th^ frames on Figure 6D right bottom corner) during which the worm changes direction (in red). We note that the omega turn appears as a combination of pauses and directed transport, highlighting that the interpretation of the classification requires *a priori* knowledge of the sample or process studied. With these four diverse examples, we have shown that wMSDc is an accessible and reliable method to distinguish directed and diffusive transport not only within the short and noisy trajectories commonly obtained in single-molecule fluorescence microscopy at the sub-cellular scale but also in various other types of trajectories occurring at different time and length scales, from single molecules, to cells and complete organisms.

## Discussion

Imaging and tracking of single biomolecules, cells and multicellular organisms has become routine over the last decade. Consequently, massive amounts of data (trajectories) have been generated, and now developing algorithms to extract quantitative information from these trajectories has become important. Most of the existing algorithms are developed using idealized, simulated trajectories: long (a large amount of time points), showing only one well-specified mode of motility (sub-diffusion, free diffusion or directed motion) and/or with low levels of noise (10, 27, 43). In practice, however, such trajectories are rarely observed, in particular for single proteins *in vivo*. Under these conditions, trajectories are often short (seconds) and noisy. Furthermore, the mode of motility can change over time or as a function of location, as is for example the case for our measurements of motor proteins moving in the chemosensory cilia of *C. elegans* (20–22, 44). To extract quantitative information from such trajectories, we developed a new method based on Mean-Square Displacements (MSD), wMSDc. We also implemented sub-classification to (i) distinguish anterograde and retrograde directed transport, and (ii) classify the type of pausing to eventually compare their durations and amplitude of displacement. Furthermore, we developed tools to extract location-specific velocities and diffusion coefficients. By analyzing simulated data (with quality comparable to our experimental data), we showed that wMSDc is a reliable approach to extract quantitative information from short and noisy trajectories. The existing windowed MSD approaches require many inputs from the user, for example for window width and threshold used to distinguish the different modes of motility. Furthermore, it has been reported that the power of the time lag (*α*) depends on the window width used (12) and the level of noise (45). Therefore, in this study, we focused on finding objective rules to select these parameters, such that the decisions that need to be taken by the user are minimal. Using simulated trajectories similar to the ones obtained experimentally for single proteins in *C. elegans*, we compared the classification accuracy of wMSDc with a Hidden Markov Model based algorithm (12) (HMM) as well as a Moment Scaling Spectrum based method (DC-MSS (13)). The comparison shows that, in particular for directed motion, the classification accuracy of wMSDc is higher in the short and relatively noisy trajectories typically observed *in vivo*.

Further development of wMSDc will be needed to use it to distinguish sub-diffusive and diffusive transport, for example by implementing a locally varying threshold (instead of a constant). wMSDc could also be combined with other state-of the art algorithms to identify the mechanisms underlying sub-diffusive and diffusive transport (e.g. Fractional Brownian motion, Continuous-Time Random Walk (29), Brownian motion). In the future, wMSDc and DC-MSS could also be combined to optimize the detection of short pauses between two sequences of directed transport, for examples employing DC-MSS to detect the short pauses (in order to limit the effects of windowing in wMSDc), and wMSDc for classification (to avoid misclassifications due to the overlap of the moment-scaling-spectrum values). wMSDc could also be combined with back-propagation neuronal networks (38) to determine the window width in order to perform classifications with a better accuracy at low velocities.

As a final demonstration of the capabilities of wMSDc, we analyzed trajectories from different experimental systems: individual motor proteins moving in the chemosensory cilia of *C. elegans*; the protein PICH moving along and interacting with double-stranded DNA held, *in vitro*, between two optical traps; migrating cancerous cells (MDA-MB-231) moving on a surface between physical traps; and *C. elegans* moving on an agar plate. The results of these analyses show that wMSDc can be used to reliably analyze a wide variety of systems, from single-molecule to whole-organism levels, in short and noisy trajectories as well as in longer and less noisy ones.

## Materials and Methods

### Simulated trajectories parameters used to compare wMSDc, HMM and DC-MSS classification efficiency

In order to set an appropriate pair (*W*, *α*_*Th*_), we focused our work on switches between direct and diffusive transports assuming that is the hardest type of trajectory to classify. The “*in-silico*” molecules mimic the motion characteristics of IFT-dynein moving in the chemosensory cilia of C. elegans (20). In this paper, we have previously quantified the duration of the pauses, the velocities, the diffusion coefficients during the pauses, and the probability of turnarounds all location-dependent. In the current simulation, the cilia is modeled by a 2D rectangle with at length of 7 μm and a width of 200 nm. The initial positions of the “*in-silico*” molecules are distributed randomly between 1 μm and 6 μm. The average duration of the trajectories equals 7 seconds in order to reproduce experimental data. The duration of the pauses varies between 1.6 s and 4.4 s depending on the location. The probability of turnarounds is also location-dependent and the probability to pause at the turnaround location is constant and equals 64%. The diffusion coefficient and velocity are location-dependent. All the parameters are summarized in Figure S2. In the Y direction, the diffusion coefficient is arbitrary set at values 3 times lower than the diffusion coefficient values in the X direction. The precision of localization is added for each X-Y positions using a Gaussian distribution centered on the precision of localization value. We simulated 9 datasets using different localization precision (*PL* = 0, 30 or 60 nm) and integration time (Δ*t* = 30, 60 or 100 ms).

A supplemental dataset (20 trajectories, *PL* = 30 nm and Δ*t* = 30 ms) combining confined diffusion (first 100 frames) and free diffusion (last 100 frames) has been generated to develop the methodology and distinguish whether the trajectories contained bouts of directed transport. Diffusion coefficient equals 0.05 *μm*^2^. *s*^−1^ for the half of the trajectories and 0.15 *μm*^2^. *s*^−1^ for the other half which is in the range of values obtained for the motor proteins (20) and transmembrane protein (46) in the chemosensory cilia of *C. elegans*. The radius of confinement during confined motion is set to 0.1 *μm*.

## Software availability

The wMSDc method and standalone application is written in Matlab and is available on GitHub (https://github.com/nbdan01/wMSDc). A dataset containing simulated and various experimental data is provided with the algorithm. The format of the dataset can be either an .xlsx file, collection of .mat files obtained with the software Fiesta or Matlab files. Data can be either 1D or 2D. The templates for the .xlsx format are also available on Github.

## Author Contributions

N.D. and E.J.G.P. designed research; N.D. performed research and analyzed data; N.D., Z.Z. and E.J.G.P. wrote the manuscript.

## Acknowledgements

We thank Andreas Biebricher, Chase Broedersz and Christine W. Bruggeman for providing the data used in the applications section (PICH proteins, MDA-MB-231 cells and *C. elegans* omega-turn data). We also thank the members of our “worm team” (Jaap van Krugten, Aniruddha Mitra, Elizaveta Loseva, Wouter Mul and Christine Bruggeman) for discussions, and in particular Jaap van Krugten for providing the first datasets that indicated a new methodology was needed. We acknowledge financial support from the European Research Council under the European Union’s Horizon 2020 research and innovation program [E.J.G.P.; Grant Agreement No. 788363 “How intraflagellar transport shapes the cilium: a single-molecule systems study” (HITSCIL)].

