## Supplemental Information for "Classifying directed and diffusive transport in short, noisy single-molecule trajectories with wMSD"

### Supplemental Figures

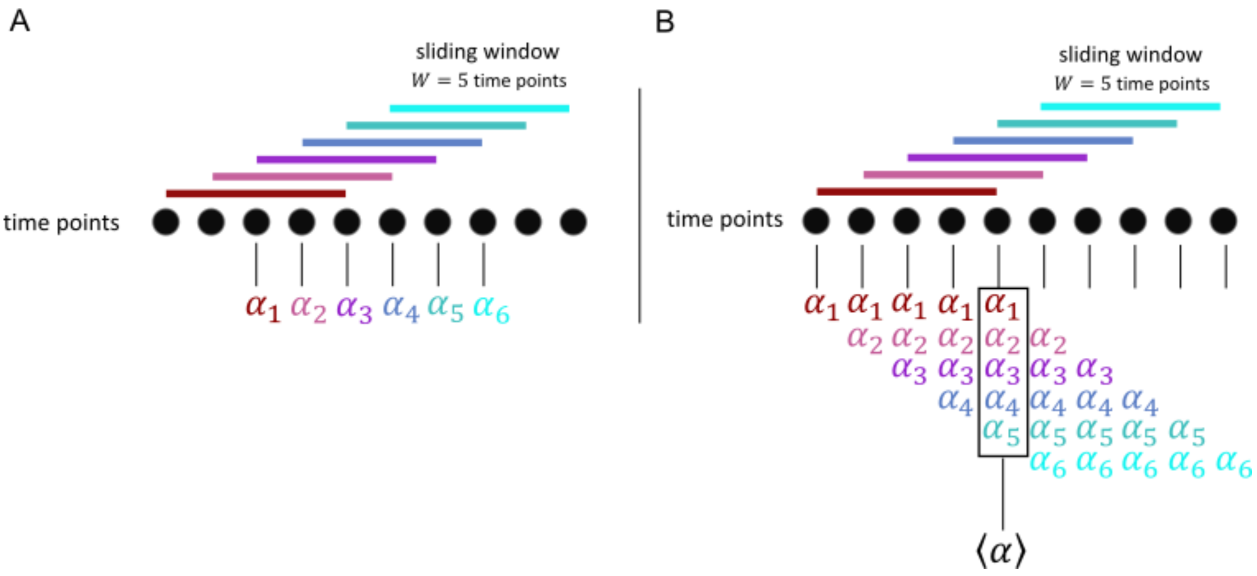


**Figure S1: Cartoon showing the method used to address a single alpha value for each time point.** In these cartoons, the window width ($W$) equals 5. A) Cartoon showing that the first $(W-1)/2^{\text{th}}$ and last $(W-1)/2^{\text{th}}$ points have not $\alpha$ value when the $\alpha_i$ value calculated within the window $W_i$ is solely addressed to the central time point of the window (where $i = 1,2,3,4 \ldots$ corresponds to the $i^{\text{th}}$ value/window). B) Cartoon showing how to address an averaged $\alpha$ value ($\langle\alpha\rangle$) for each time point in the trajectory. The $\alpha_i$ value calculated within each window $W_i$ is addressed for all the time points included in $W_i$. In this way, each points have one or more $\alpha_i$ values. The final $\alpha$ value for each point corresponds to the average of all the $\alpha_i$ values ($\langle\alpha\rangle$).

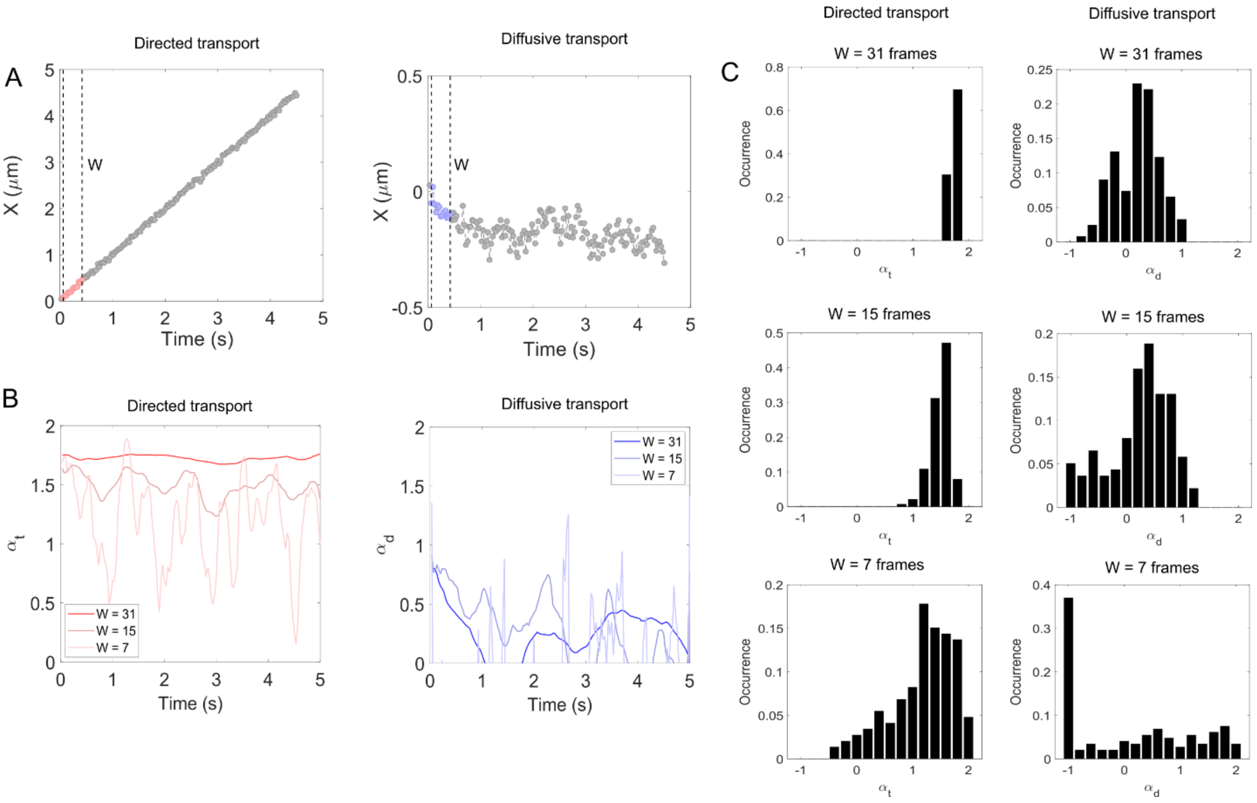


**Figure S2: Distribution of alpha values depending on the Window width used.** A) X-T (1D) simulated trajectories for directed ($v$ = 1 $\mu m.s^{-1}$, $\Delta t$ = 30 ms and $PL$ = 30 nm) and diffusive ($D$ = 0.01 $\mu m^2.s^{-1}$, $\Delta t$ = 30 ms and $PL$ = 30 nm) transport. The window width (W) equals 15 frames. B) Fluctuations of alpha values over time for directed (left) and diffusive (right) transport: the narrower the window width, the higher the amplitude of the fluctuations. These trajectories are also used in Figure 1. C) Histograms of alpha values calculated for directed (left) and diffusive (right) transport using different window widths (W). The narrower the window width, the wider the spread of the alpha values, due to fluctuations of alpha on short time scale.

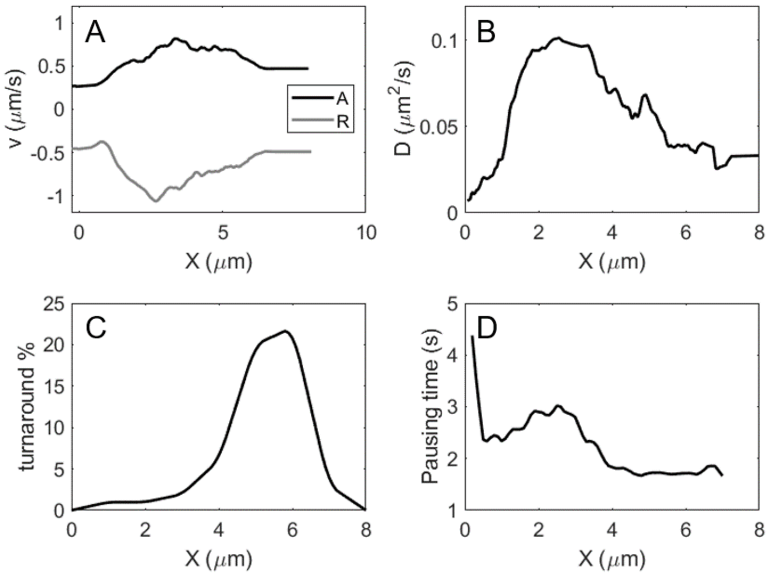


**Figure S3: Parameters used to generate turnaround trajectories.** A) Velocities in the anterograde and retrograde direction, B) Diffusion coefficient, C) Percentage of turnarounds, D) Pausing time when a turnaround occurs. These values mimic the motion characteristics of IFT-dynein moving in the chemosensory cilia of *C. elegans* (18). See also Methods section.

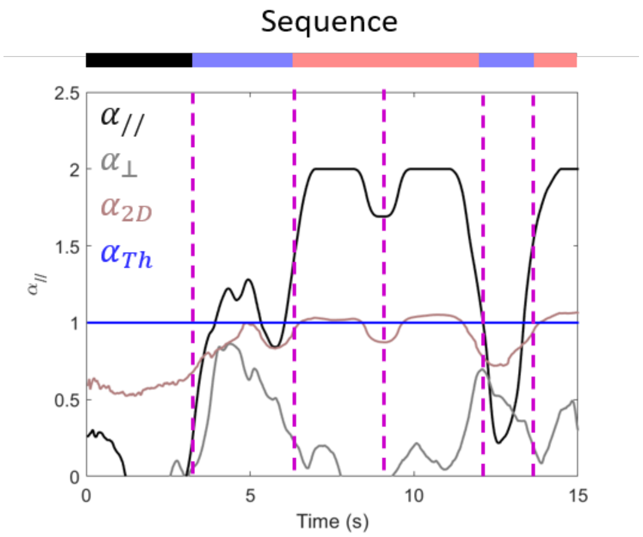


**Figure S4: Alpha values for a simulated trajectory containing sub-diffusion, free diffusion and directed motion.** Alpha values in the parallel (//) and perpendicular (⊥) direction (black and grey curves) as well as alpha values calculated using the MSD in 2 dimensions (pink curve). The alpha threshold determined using Eq.3 and Condition #1 is shown in dark blue. The dotted purple vertical lines show the separation between the different types of motion. The complete sequence is shown on the top: Dark: sub-diffusion, Blue: free diffusion, Red: directed motion. The first directed motion sequence (in red) corresponds to a direct turnaround (anterograde-directed transport followed by retrograde-directed transport). The dotted vertical line in the middle indicates the location of the turnaround.

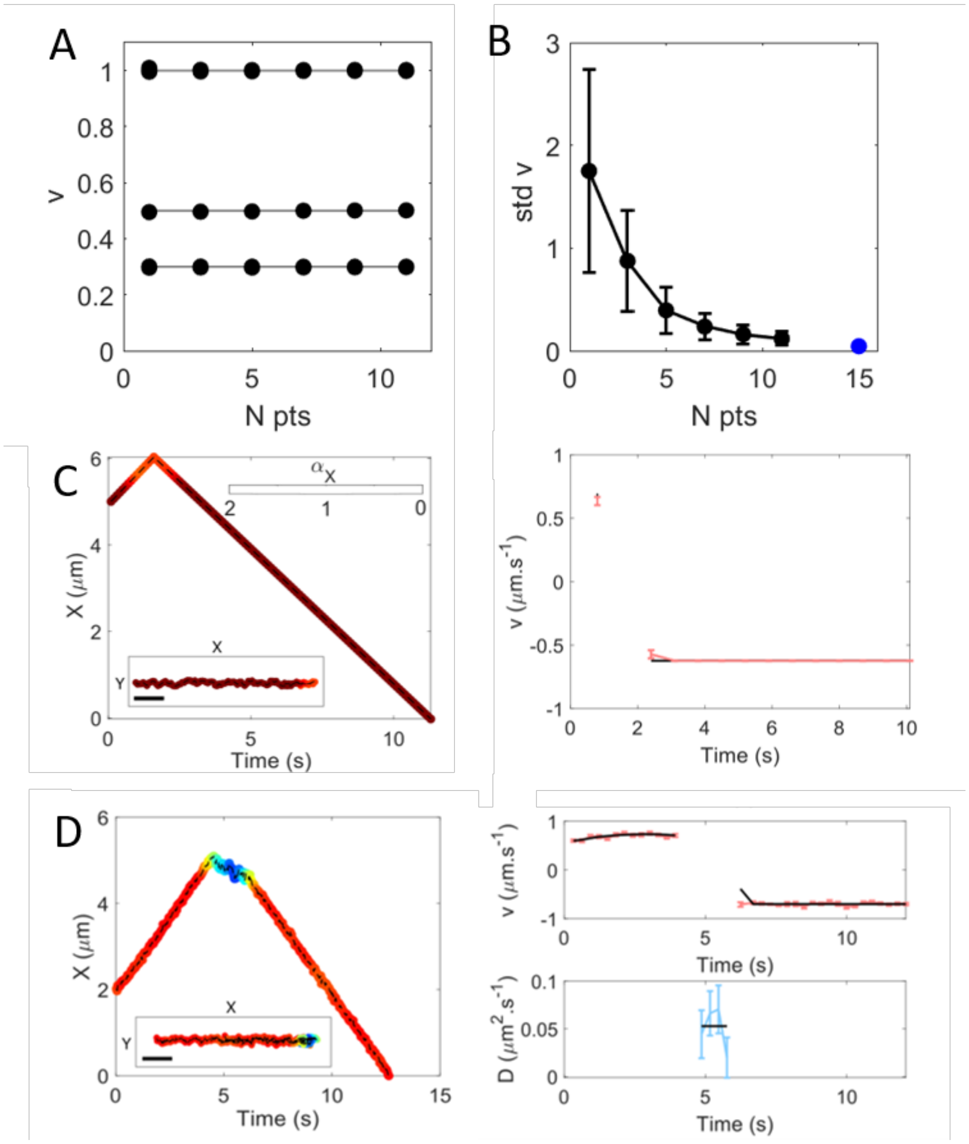


**Figure S5: Average velocities as a function of the number of time points used to calculate them.** A) Average velocity (black circles) obtained using our approach. The black lines indicate the velocities injected in the simulation. The parameters of the simulated data are listed in the Supplementary Table 1 (Set T). B) Standard deviation of the mean as a function of the number of time points used to calculate the velocity (black circles). The blue circle corresponds to the standard deviation of the mean obtained using a windowed Mean-Square Displacement over 15 frames. C,D) Examples showing the velocities and diffusion coefficients obtained for two simulated trajectories (those from Figure 4 A and B). The black lines correspond to the input values. The red and blue error bars show the location/time dependent velocities and diffusion coefficients respectively (mean and standard deviation of the mean calculated over 8 points).

### Supplemental Tables

| # Set D | D (um^2/s) | PI (um) | dt (s) |
|---|---|---|---|
| 1 | 0.005 | 0.03 | 0.03 |
| 2 | 0.01 | 0.03 | 0.03 |
| 3 | 0.02 | 0.06 | 0.03 |
| 4 | 0.025 | 0.03 | 0.03 |
| 5 | 0.035 | 0.06 | 0.03 |
| 6 | 0.09 | 0.03 | 0.03 |
| 7 | 0.09 | 0.06 | 0.03 |
| 8 | 0.1 | 0.03 | 0.03 |
| 9 | 0.1 | 0.03 | 0.06 |
| 10 | 0.1 | 0.03 | 0.09 |
| 11 | 0.1 | 0.06 | 0.03 |
| 12 | 0.1 | 0.06 | 0.06 |
| 13 | 0.1 | 0.06 | 0.09 |
| 14 | 0.35 | 0.06 | 0.03 |
| 15 | 0.45 | 0 | 0.03 |
| 16 | 0.45 | 0.03 | 0.03 |
| 17 | 1.8 | 0.06 | 0.03 |
| 18 | 1.8 | 0.06 | 0.03 |
| 19 | 1.8 | 0.06 | 0.09 |
| 20 | 1.8 | 0.06 | 0.03 |
| 21 | 1.8 | 0.06 | 0.05 |
| 22 | 1.8 | 0.06 | 0.03 |
| 23 | 1.8 | 0.06 | 0.09 |
| 24 | 1.8 | 0.09 | 0.09 |

| # Set T | v (um/s) | PI (um) | dt (s) |
|---|---|---|---|
| 1 | 0.3 | 0.03 | 0.03 |
| 2 | 0.3 | 0.03 | 0.06 |
| 3 | 0.3 | 0.03 | 0.09 |
| 4 | 0.3 | 0.06 | 0.03 |
| 5 | 0.30 | 0.06 | 0.06 |
| 6 | 0.3 | 0.06 | 0.09 |
| 7 | 0.30 | 0.06 | 0.09 |
| 8 | 0.5 | 0.06 | 0.03 |
| 9 | 0.75 | 0.06 | 0.03 |
| 10 | 0.8 | 0.06 | 0.05 |
| 11 | 1 | 0 | 0.03 |
| 12 | 1 | 0.03 | 0.03 |
| 13 | 1 | 0.03 | 0.03 |
| 14 | 1 | 0.03 | 0.03 |
| 15 | 1 | 0.03 | 0.03 |
| 16 | 1 | 0.03 | 0.03 |
| 17 | 1 | 0.06 | 0.03 |
| 18 | 1 | 0.06 | 0.03 |
| 19 | 1 | 0.06 | 0.03 |
| 20 | 1 | 0.06 | 0.03 |
| 21 | 1 | 0.06 | 0.03 |
| 22 | 1 | 0.06 | 0.03 |
| 23 | 1 | 0.06 | 0.09 |
| 24 | 1 | 0.09 | 0.09 |

**Table S1: Set D and T used to characterize the MSD analysis outputs values for passive and directed transport and taking in account the integration time and Precision Inaccuracy.** Values of the diffusion coefficients (D), velocities (v), integration times (dt) and Precision of localisation (PL) used to characterize the MSD analysis outputs values.

| Type of data | $\langle v \rangle$ ($\mu m. s^{-1}$) | $v_{min}$ ($\mu m. s^{-1}$) | Smoothing for $N_{A/R}$ | Settings |
|---|---|---|---|---|
| Motor proteins *in vivo* ($\Delta t$ = 30-50 ms) | 0.6 | 0.2 | 30 | Default settings |
| Motor proteins in dendrite ($\Delta t$ = 150 ms) | 0.6 | 0.3 | 30 | $\alpha$ threshold fixed at 1.3-1.4 or Default Setting. |
| PICH protein *in vitro* ($\Delta t$ = 1-2.5 s) | 0.025 | 0.01 | 30 | Default settings<br>Decrease at 30%<br>1D data<br>Condition #2,3 and 4 removed |
| Cancer cells ($\Delta t$ = 10 minutes) | 0.01 | 0.0025 | 15 | Advanced settings<br>Decrease at 30%<br>Eq. 3bis (using the median) |
| C. elegans ($\Delta t$ = 1 frame) | 1.2 | 1.6 | 30 | Default settings |
| Confined and Free diffusion (simulation) | 0.6 | 0.1 (D = 0.15 $\mu m^2. s^{-1}$)<br>0.3 (D = 0.05 $\mu m^2. s^{-1}$) | 30 | Default settings |
| Directed and diffusive transport (simulated) | 0.8 | 0.7 | 30 | Default settings |

**Table S2: Input parameters used for the classification with wMSDc.** Values of the different parameters and conditions used to perform the classification on experimental data. $\langle v \rangle$ and $v_{min}$ correspond to the estimated average and minimal velocities ($\mu m \cdot s^{-1}$). These parameters are used to determine the appropriate window width. $N_{A/R}$ corresponds to the number of time points used to perform the sub-classification between anterograde and retrograde directed transport. The 4$^{th}$ column corresponds to the settings used to determine the threshold in order to perform the classification between diffusive and directed transport.
